# Nuclease-Resistant L-DNA Tension Probes Enable Long-Term Force Mapping of Single Cells and Cell Consortia

**DOI:** 10.1101/2024.07.22.604597

**Authors:** Soumya Sethi, Tao Xu, Aritra Sarkar, Christoph Drees, Claire Jacob, Andreas Walther

## Abstract

DNA-based tension probes with precisely programmable force response provide important insights into cellular mechanosensing. However, their degradability in cell culture limits their use for long-term imaging, for instance, when cells migrate, divide, and differentiate. This is a critical limitation for providing insights into mechanobiology for these longer-term processes. Here, we present DNA-based tension probes that are entirely designed based on the stereoisomer of biological D-DNA, i.e., L-DNA. We demonstrate that L-DNA tension probes are essentially indestructible by nucleases and provide days-long imaging without significant loss in image quality. We also show their superiority already for short imaging times commonly used for classical D-DNA tension probes. We showcase the potential of these resilient probes to image minute movements, and for generating long term force maps of single cells and for the first time, of collectively migrating cell populations.

---

Cells constantly sense and respond to a vast number of physical stimuli within the extracellular matrix. Mechanotransduction is the process whereby mechanical signals are transduced into biochemical signals to regulate cell fate.^[1]^ Mechanotransduction pathways are mediated by interactions between cell surface receptors and the cell cytoskeleton. Numerous cell surface receptors such as integrins,^[2]^ cadherins,^[3]^ notch,^[4,5]^ tyrosine kinases^[6]^ etc. bidirectionally transmit forces to their respective ligands on the extracellular matrix. Such receptor-ligand interactions generate piconewton forces (pN) as cells transverse the extracellular matrix.^[7]^ Mechanobiology plays an integral role in cellular processes such as cell migration, cell division, wound healing,^[8]^ stem cell lineage guidance,^[9]^ differentiation, as well as patterning and organization of germ layers during embryonic development.^[10]^ These remarkably complex and multicellular processes span over multiple days, and the role of mechanotransduction in these phenomena at a molecular level remains largely unexplored.

Over the past decade, DNA based molecular tension probes have paved the path for understanding the complexities of cellular mechanotransduction.^[7]^ Such probes typically contain a DNA duplex and a judiciously placed fluorophore and quencher pair, as well as a cell surface-receptor ligand, such as RGD for binding to integrins. Depending on the geometry, most prominently zipper, shear or peel-off,^[11]^ the force for rupture can be engineered from a few pN to ca. 70 pN.^[7]^ Cellular tension forces can thus be measured by direct appearance of fluorescence signals. The probes can either be designed for an irreversible (duplex)^[12–16]^ or reversible (hairpin)^[17–20]^ force detection, allowing to map cellular traction forces in real time. Similar probes can also be integrated into hydrogels to design mechano-reporting materials.^[21]^

Despite many advances in DNA-based tension probes, they suffer from the major drawback that typical cell culture environments contain nucleases that degrade DNA. More specifically, these enzymes degrade the biological stereoisomer of DNA, that is D-DNA. This critically limits the overall lifetime of tension probes and compromises signal quality (signal-to-noise) early on. Fetal bovine serum (FBS)– the gold standard for cell culture – contains greater than 256 U/L equivalent of DNase 1 activity,^[22]^ as well as other nucleases.^[23–25]^ FBS is an important supplement to maintain cell phenotype and cell growth, and remains hard to avoid. This nuclease activity restricts cellular force mapping to only short periods of time, and even nuclease inhibitors perform poorly, thus impeding the study of force generation in various important long-term phenomena like cell migration, division, and differentiation. Consequently, current D-DNA based probes are restricted to single cell mechanobiology studies at comparably short time scales. To delve further into grasping new realms of mechanobiology, nuclease-resistant and stable probes are needed.

Here, we introduce L-DNA mechanoprobes, based on the non-biological stereoisomer of D-DNA, that are remarkably bioinert and resistant to nucleases. These probes can sustain cell culture environments for several days without visible degradation. We also show that even during short-term measurements, presently used in mechano-profiling of cells, L-DNA presents critical advantages in imaging without extensive background subtraction. We display for the first time that DNA based probes can map forces of cell consortia. We also demonstrate their applicability for studying mechanotransduction during important cellular behaviors like membrane ruffling, cell division, and collective cell migration.

Building on the knowledge of zipper-type DNA mechanoprobes,^[7]^ we designed L-DNA tension probes that we anchored via biotin-neutravidin chemistry to surfaces (Figure 1). We specifically opted for L-DNA and not other types of locked nucleic acids^[26]^ or also peptide nucleic acids, as the stereoisomer of biological D-DNA follows exactly the same physics regarding mechano-activation as the biological D-DNA sister. Hence, forces can be predicted using known relationships, and even simulations based on D-DNA can be run based on known force fields^[27,28]^ and directly translated to our L-DNA systems (Figure 2).

**Figure 1.**
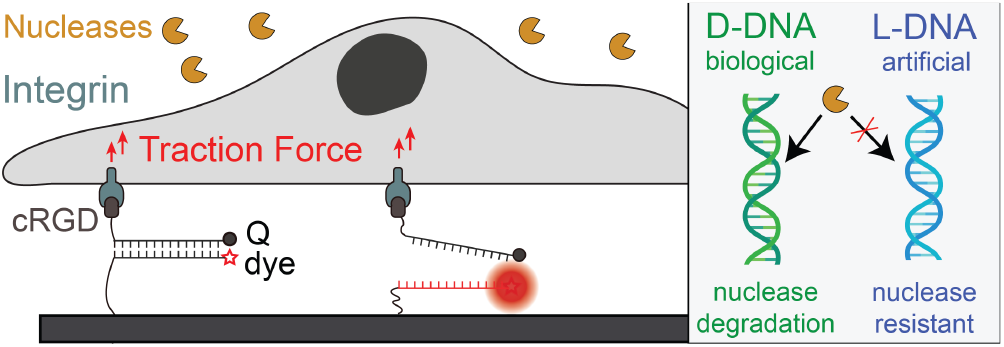
Scheme depicting DNA-based probes interacting with cells. Biological D-DNA probes degrade rapidly by nuclease action in cell culture environments whereas the non-biological stereoisomer L-DNA probes are highly resistant.

**Figure 2.**
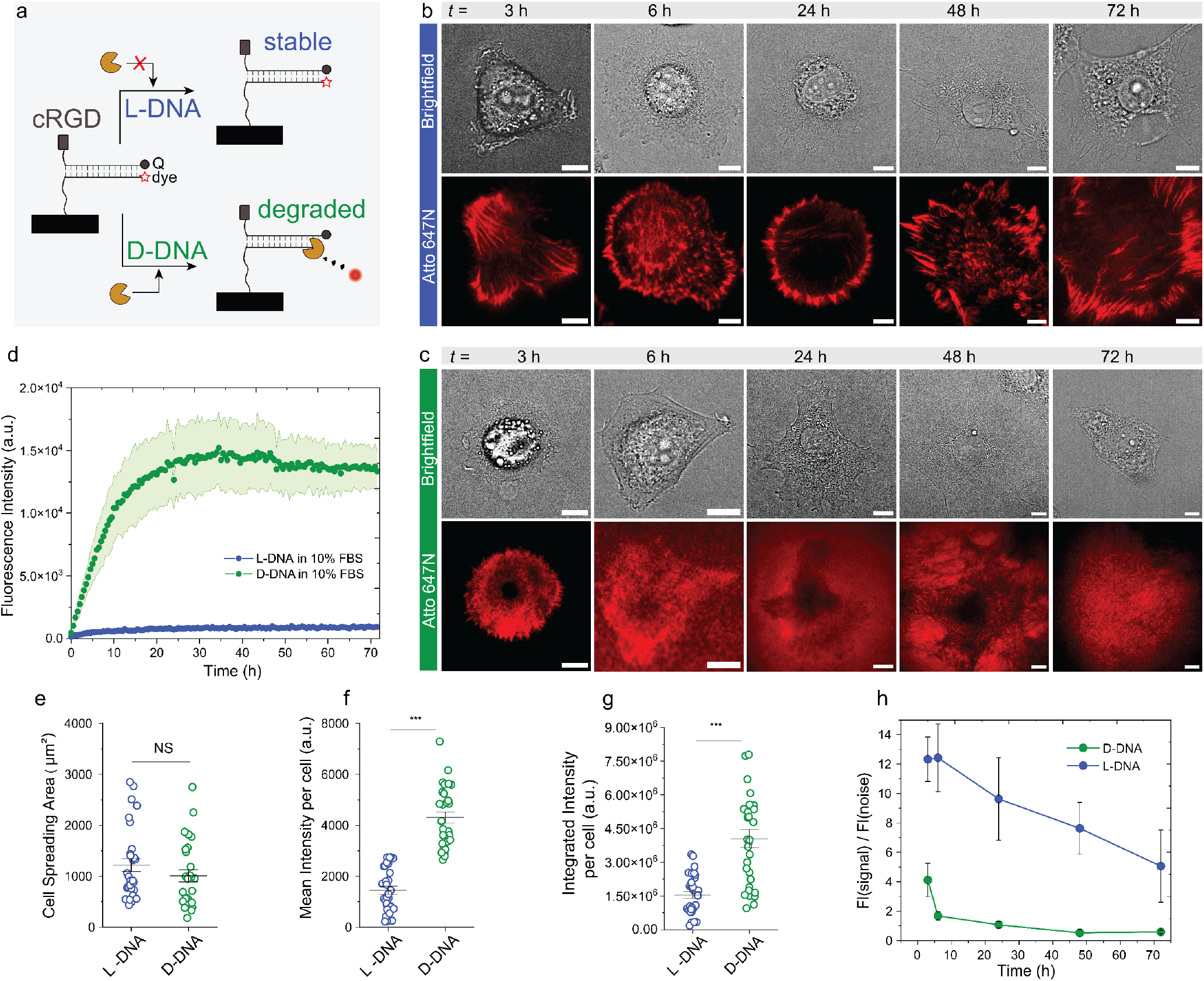
Comparison of D-DNA versus L-DNA tension probes. (a) Scheme illustrating the degradation of D-DNA and stability of L-DNA probes in the presence of nucleases present in cell culture medium. The tension probe in this case is in unzipping geometry with 12 pN unzipping force. (b-c) Brightfield and fluorescence images of fibroblasts/NIH3t3 cells on (b) L-DNA and (c) D-DNA tension probe surfaces at 3 h – 72 h after seeding. Scale bars = 10 µm. (d) Degradation kinetics of fluorophore/quencher pair-containing L-DNA and D-DNA duplexes in cell culture media at 37 °C. (e) Comparison of cell spreading area at 3 h on D-DNA and L-DNA tension probe surfaces (n = 30 cells; NS = not significant). (f) Comparison of mean fluorescence intensity on L-DNA and D-DNA tension probe surfaces at 3 h (n= 30 cells, ^***^ p<0.001), unpaired student’s t-test). (g) Comparison of integrated fluorescence intensity per cell L-DNA and D-DNA tension probe surfaces at 3 h (n = 30 cells, ^***^ p<0.001). (h) Signal to noise ratio as defined by mean fluorescence intensity (FI) per cell divided by background mean intensity (n = average of 10 cells/time point, error = standard deviation).

We first investigated the stability of L-DNA versus D-DNA probes in solution by studying the degradation kinetics of fluorophore/quencher pair-containing L-DNA and D-DNA duplexes in cell culture media containing 10% FBS at 37 °C. Figure 2d clearly depicts that D-DNA undergoes rapid degradation, as seen by the fluorescence increase within the first few hours. It is fully degraded within one day. In contrast L-DNA is highly stable for days, here measured for 72 h. We also assessed the stability of a fluorophore-containing L-DNA immobilized under typical conditions on a microscopy slide (Supplementary Figure S1). To this end, we immobilized a single stranded L-DNA dye conjugate onto a coverslip and measured the fluorescence intensity in total internal reflection microscopy (TIRF) during incubation with FBS-containing cell culture media. Indeed, the fluorescence of the surface-bound L-DNA only shows a minor fluorescence decrease during the observation period of 3 days. This confirms a very robust DNA mechanosensing platform, promising days of unimpeded observation of cellular traction forces.

Next, we turn to the direct comparison between D-DNA and L-DNA tension probe surfaces by analyzing the tension signals generated by fibroblasts over a period of 3 days in 10% FBS cell culture medium at 37 °C. Briefly, the geometry of the tension probe was designed to be in an unzipping mode^[13]^ such that the dsDNA duplex is denatured one base at a time. The force required to pull apart the duplex is around 12 pN. The tension probes were labelled with a fluorophore-quencher pair along with cRGDfk ligand, which has a high affinity to the αvβ_3_integrin receptors (Figure 1).^[29]^ After seeding cells on the D-DNA and L-DNA surfaces, we imaged them at different time points from 3 – 72 h using TIRF microscopy. Figure 2b,c depict representative single cells and Supplementary Figures S2-S6 show detailed overviews of more examples. Importantly, these images are not additionally treated by background subtraction routines. Mechanoactivation can be observed for both D-DNA and L-DNA surfaces. For the D-DNA mechanosensors, the fluorescence signals can be well detected at 3 h, but the signal quality deteriorates heavily at 6 h, and over longer times degradation and noise across the entire field of view dominate the image. In striking contrast, the fluorescence signal for L-DNA probes remains detailed and vivid even as time progressed to 72 h. Even at an imaging time of only 3 h, a crisper signal is evident in the direct comparison between L-DNA and D-DNA surfaces. Thus, even at short imaging time frames, L-DNA has a clearly superior performance. It may be noted that the image quality of the D-DNA surface can be improved by a background subtraction of the images up to 6 h, but this already contains assumptions in image treatment, such as equal degradation of probes below and adjacent to the cells (Supplementary Figure S7). Even after background subtraction, the non-background subtracted L-DNA images at 6 h are clearly more defined (compare Supplementary Figure S3 to S7). This data demonstrates the significant beneficial effect of the enhanced stability of the L-DNA probes. Imaging for 3 days is without problems.

We further quantified the cell spreading area of each cell, the mean intensity of the fluorescence signal produced by each cell, and the total amount of fluorescence signal produced by each cell on both surfaces (Figure 2e-g). At 3 h, the cell spreading areas on both surfaces is comparable, whereas the mean intensity and the integrated fluorescence signal per cell is significantly higher in the case of D-DNA probes. We submit that this effect rather stems from degradation of the D-DNA probes as early as 3 h after seeding cells (see degradation kinetics in Figure 2d), thus leading to a contamination of true mechano-signaling. Due to the nuclease resistance of the non-biological L-DNA probes, the L-DNA surfaces report more accurately the true mechano-signaling events.

Due to the concurrent degradation, a further evaluation of these three parameters at longer time frames does not make sense. Therefore, we set out to compare the mean of the fluorescence intensity signal per cell to the background mean fluorescence intensity adjacent to the cell, termed as signal-to-noise ratio, over a 3-day time frame (Figure 2h). A fivefold higher signal-to-noise ratio in favor of L-DNA is already visible at 3 h, indicating significant contamination of the cell traction force signals by degradation-induced fluorescence in D-DNA. As time progresses, the D-DNA signal-to-noise ratio plummets falling below 1 at the 24 h time point, indicating that the noise fluorescence intensity surpasses the signal fluorescence intensity. In comparison, the signal-to-noise ratio of the L-DNA probe remains relatively constant from 3 to 6 h and then progressively decreases. We suspect this to occur because cell migration and proliferation produce tension signals in adjacent areas during longer imaging times, thereby increasing the background fluorescence. At 6 h, the signal-to-noise ratio of the L-DNA probes is 10 times higher than for D-DNA probes. This is a significant advantage for long term imaging. Supplementary Video 1 underscores the differences.

Encouraged by the stability of duplex L-DNA probes, we embarked on using reversible DNA hairpin probes to understand how imaging of even minute movements of cells in cell culture environments particularly challenging for DNA could benefit from nuclease-resistant probes. Building on the work by Salaita and co-workers,^[17]^ we designed D-DNA and L-DNA hairpin probes that can reversibly map cell forces in the range of around 16 pN (Supplementary Table S1). We opted for a myoblast cell line (A10), that requires culturing conditions at 20% FBS, and which has the capability of membrane ruffling as they migrate on surfaces. Figure 3 depicts a direct comparison. We imaged the cells 18 h after seeding them on DNA coated surfaces. For surfaces coated with D-DNA, the tension signals are difficult to locate (Figure 3c) owing to the rapid degradation of these D-DNA probes in 20% FBS. Even though the brightfield images show some level of membrane ruffling, the corresponding fluorescence image does not show significant changes in the cross-sectional analysis (Figure 3d). The surroundings of the cell are equally bright as the cell edge where membrane ruffling takes place. In contrast, the L-DNA surfaces exhibit clear and detailed signals that reversibly appear and disappear as the membranes ruffle (Supplementary Video 2). While we are not saying that the membrane ruffling may not be observable at all using D-DNA probes, e.g., by heavy optimization of culture conditions or time frames, we emphasize that the L-DNA hairpin probes allow for a straightforward and clear imaging of even such minute movements without heavy alterations of the culture conditions that may be detrimental to normal cell behavior. These results underscore that L-DNA probes can open new avenues for our understanding of mechanobiology, for instance to map forces for cells that require longer to adhere onto surfaces.

**Figure 3.**
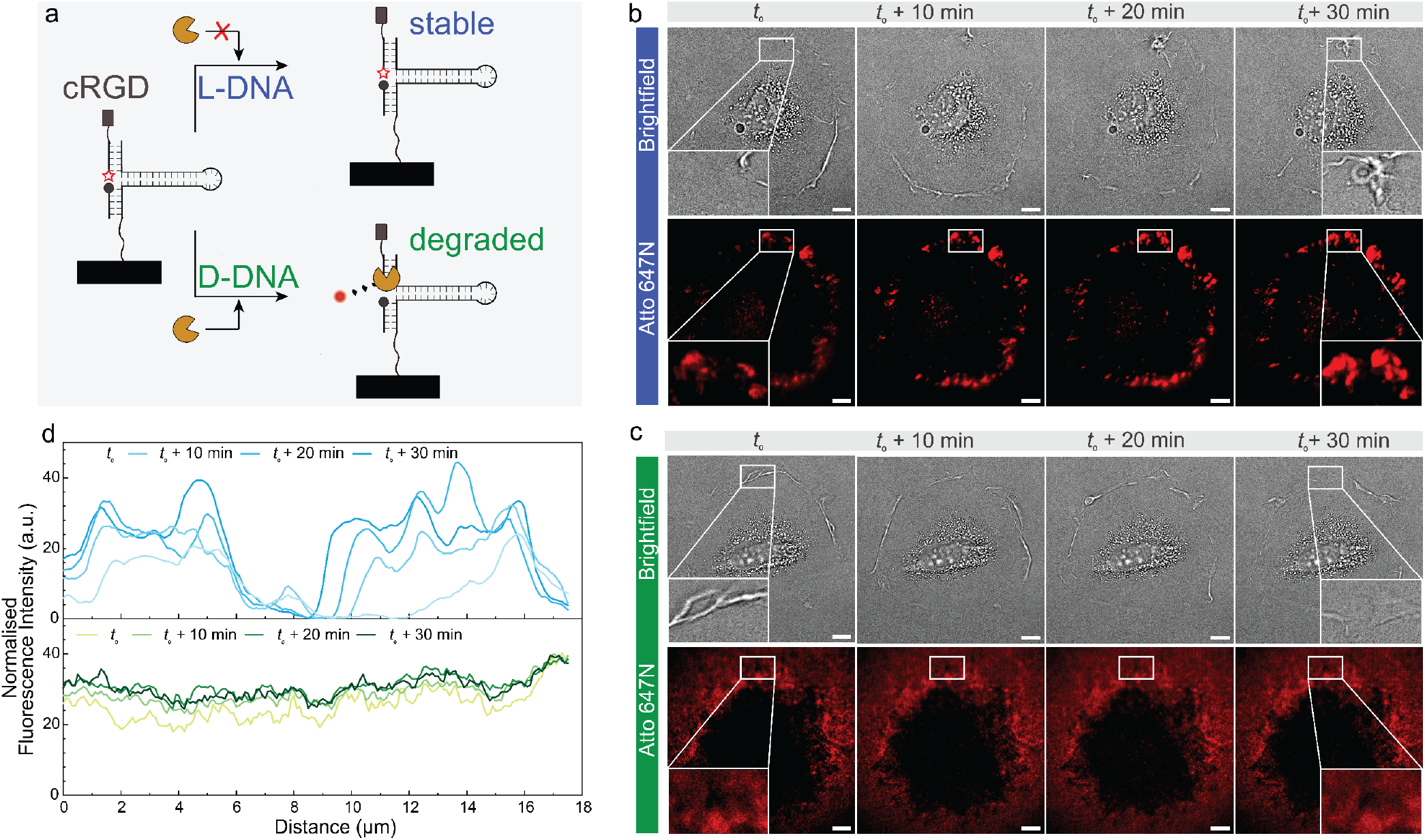
Comparison of D-DNA and L-DNA reversible hairpin probes to study membrane ruffling in myoblasts (A10). (a) Schematic depicting the geometry of hairpin tension probes and the stability in cell culture media. The used hairpin probes have 16 pN opening force. (b-c) Time lapse brightfield and fluorescence images on (b) L-DNA and (c) D-DNA hairpin tension probe surfaces 18 h after cell seeding. Consecutive images depict the ruffling of the myoblast cell membrane as illustrated by a cross-sectional region of interest (ROI). Scale bar = 10 µm, (d) Cross sectional horizontal analysis of the fluorescence intensity at different time points within the ROI from b and c depicting ruffling of the cell membrane as the cell migrates forward.

Finally, we tackled the challenge of going beyond imaging a single cell and investigated force mapping during collective cell migration. The term collective migration of cells implies migration of cells as a unified group in sheets and clusters. *In vivo*, collective cell migration is the hallmark of re-modelling events such as morphogenesis, wound repair, and cancer invasion.^[30]^ *In vitro*, the exclusion method and the scratch method are important assay techniques. The exclusion method involves seeding cells at a high density on two sides of a barrier, culturing them until confluency and then removing the barrier to allow cells to migrate. The scratch method involves manual scratching of a cellular monolayer to simulate a wound. We opted for the exclusion method due to its compatibility with surface modifications. On surfaces covered with duplex tension probes in unzipping geometry, we positioned a physical barrier to confine cell seeding in two compartments. After seeding fibroblasts, we allowed them to settle and spread for about 3 h, *t*_0_, and then removed the barrier, and monitored their migration across the surfaces.

As the cells migrate in unison as sheets on L-DNA surfaces, comprehensive and intricate force maps can be observed at every time point (Figure 4b, Supplementary Video 3). In contrast, on D-DNA surfaces, only background noise can be detected at different time points (Figure 4c). Fluorescence line intensity profiles underscore the differences (Figure 4d). For the L-DNA tension probe samples, the fluorescence intensity increases over time in the barrier region, whereas D-DNA tension probe surfaces display no significant changes and high background due to excessive degradation. Consequently, L-DNA probes can effectively map forces as cells migrate cohesively and collectively. This finding opens new avenues for mechanobiology studies, as understanding of molecular-scale force mapping can give new insights into various molecular mechanisms and pathways during complex biological processes such as wound healing and morphogenesis.

**Figure 4.**
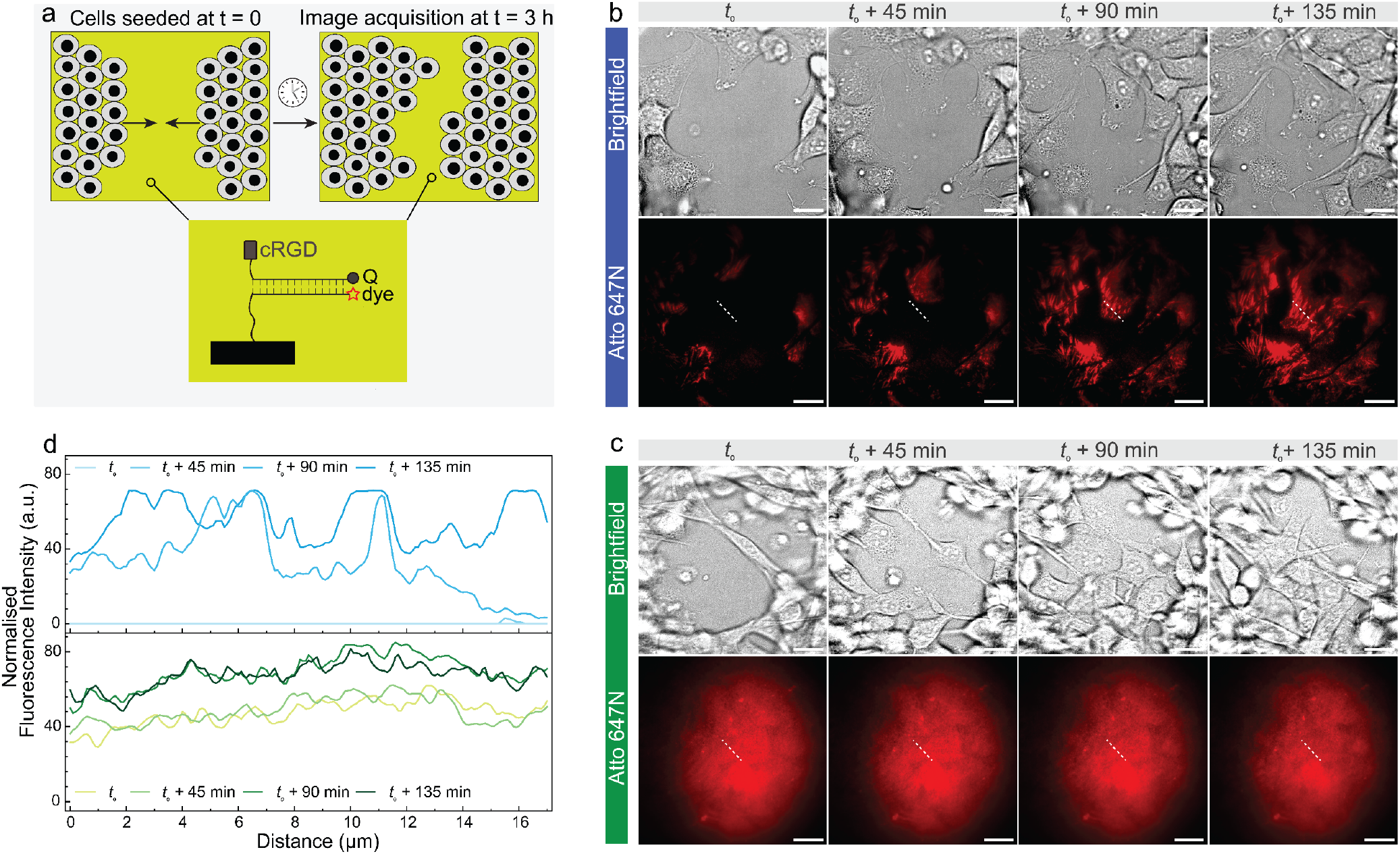
Mapping forces during collective cell migration. (a) Schematic representation (b-c) Time lapse Brightfield and fluorescence images as fibroblasts migrate collectively on (b) L-DNA and (c) D-DNA tension probe-coated surfaces. Captured 3 h after seeding, and removal of a 500 µm barrier separating two fibroblast sheets. Scale bars = 20 µm. (d) Overlay of selected intensity cross sections from b and c.

In summary, we have introduced L-DNA – the non-biological stereoisomer of D-DNA – as a powerful and highly durable cellular mechanoprobe, enabling detailed traction force mapping of single cells as well as for collective cell behavior. In direct comparison, L-DNA probes show drastically reduced degradation in classical cell culture medium compared to D-DNA probes, and thus enable long term imaging of cellular traction forces. D-DNA probes break down in a time frame of 6 h, and are already clearly compromised at 3 h, whereas L-DNA probes can be imaged for days without significant loss in quality. The signal-to-noise ratio is 5 to 10 times higher even at early time points. This is due to the impossibility of nucleases to degrade L-DNA. The advantages manifest in particular when targeting cellular behavior that requires extended culture times before starting the image acquisition of long-term behavior such as collective migration.

In contrast to other approaches for improved nuclease-stability, such as chemically altered nucleoside analogs like locked nucleic acids, 2’-O-methyl ribonucleotides, phosphorothioate linkages, or peptide nucleic acids, which hamper or change the hybridization properties and alter the force response in an unknown fashion, L-DNA offers complete biostability along with identical physical properties, hybridization kinetics, thermal stability, and force behavior compared to the biological D-DNA.^[31]^ This allows for a very simple adaptation. Further progress supports the use of L-DNA. L-DNA or L-RNA aptamers can be selected as so-called spiegelmers^[32,33]^ against the enantiomer of the target ligand, thereby having a high degree of specificity along with bio-inertness and non-immunogenic behavior. Such L-DNA/L-RNA aptamers are known for small molecules, proteins, and peptides^[34,35]^ and are in clinical trials.^[36]^ Recent studies have also shown that the L-DNA and D-DNA worlds can be connected in strand displacement reactions using peptide nucleic acids, thus substantially extending possibilities in applications.^[37]^ Additionally, D-amino acid versions of DNA polymerases and RNA polymerases have been shown to amplify L-RNA and transcribe L-DNA into L-RNA respectively.^[38–40]^ Despite still being in its infancy, the field of mirror image oligonucleotides holds immense potential. Looking to the future, the use of L-DNA mechanosensors opens new avenues for more accurate short term mechanobiology investigations, and critically enables long term imaging. The emerging concepts surrounding synthetic manipulation of L-DNA, and the combination with the D-DNA world provide ample opportunities to design robust, bioinert, and non-immunogenic downstream reactions of the mechano-activated cryptic DNA sites for building intelligent mechano-interfaces to cells.

## Supporting Information

The authors have cited additional references within the Supporting Information.^[16]^

Supporting Information File 1. Supporting Video Files 1-3.

## Acknowledgements

This project has received funding from the European Research Council (ERC) under the European Union’s Horizon 2020 research and innovation program (M3ALI: 101001638), as well as from the CoM2Life start-up funds of JGU. T.X. acknowledges support from the Max Planck Graduate Center. A.S. acknowledges funding from the Alexander von Humboldt Fundation. AW acknowledges funding via a Gutenberg Research Professorship underpinning his Life-Like Materials Program. The Zeiss Elyra 7 was funded by the Gutenberg Research College and Deutsche Forschungsgemeinschaft (DFG; 497845157).

## Entry for the Table of Contents

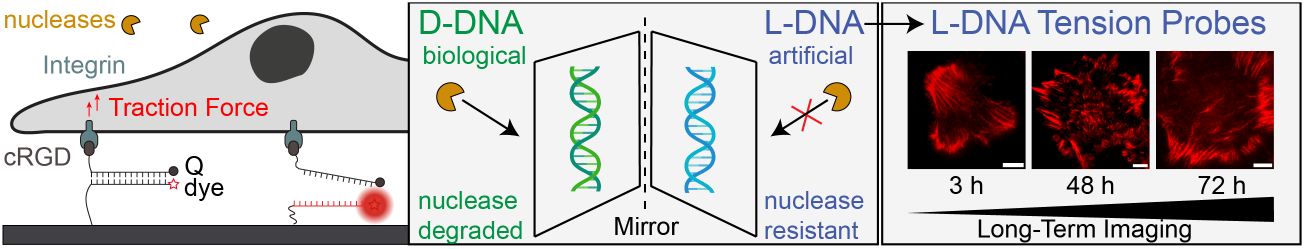

We present a novel class of DNA-based tension probes, L-DNA tension probes, based on the non-biological stereoisomer of D-DNA. These probes are exceptionally resistant to nuclease degradation and remarkably bioinert. Unlike conventional D-DNA probes, L-DNA probes can endure cell culture environments for several days, enabling long-term imaging of single cells as well as a group of cells.

Institute and/or researcher Twitter usernames: @WaltherLab, @soumyasethi94

## References

[1] D. E. Ingber, FASEB J. Off. Publ. Fed. Am. Soc. Exp. Biol. 2006, 20, 811–827.

[2] M. A. Schwartz, Cold Spring Harb. Perspect. Biol. 2010, 2, a005066.

[3] N. Borghi, M. Sorokina, O. G. Shcherbakova, W. I. Weis, B. L. Pruitt, W.J. Nelson, A. R. Dunn, Proc. Natl. Acad. Sci. U. S. A. 2012, 109, 12568– 12573.

[4] V. C. Luca, B. C. Kim, C. Ge, S. Kakuda, D. Wu, M. Roein-Peikar, R. S. Haltiwanger, C. Zhu, T. Ha, K. C. Garcia, Science 2017, 355, 1320– 1324.

[5] Y. Narui, K. Salaita, Biophys. J. 2013, 105, 2655–2665.

[6] K. Salaita, P. M. Nair, R. S. Petit, R. M. Neve, D. Das, J. W. Gray, J. T. Groves, Science 2010, 327, 1380–1385.

[7] Y. Liu, K. Galior, V. P.-Y. Ma, K. Salaita, Acc. Chem. Res. 2017, 50, 2915–2924.

[8] B. Kuehlmann, C. A. Bonham, I. Zucal, L. Prantl, G. C. Gurtner, J. Clin. Med. 2020, 9, 1423.

[9] C. Ferrai, C. Schulte, Eur. J. Cell Biol. 2024, 103, 151417.

[10] Y.-C. Poh, J. Chen, Y. Hong, H. Yi, S. Zhang, J. Chen, D. C. Wu, L. Wang, Q. Jia, R. Singh, W. Yao, Y. Tan, A. Tajik, T. S. Tanaka, N. Wang, Nat. Commun. 2014, 5, 4000.

[11] R. Ma, S. A. Rashid, A. Velusamy, B. R. Deal, W. Chen, B. Petrich, R. Li, K. Salaita, Nat. Methods 2023, 20, 1666–1671.

[12] X. Wang, T. Ha, Science 2013, 340, 991–994.

[13] Y. Duan, R. Glazier, A. Bazrafshan, Y. Hu, S. A. Rashid, B. G. Petrich, Y. Ke, K. Salaita, Angew. Chem. Int. Ed. 2021, 60, 19974–19981.

[14] J. M. Brockman, H. Su, A. T. Blanchard, Y. Duan, T. Meyer, M. E. Quach, R. Glazier, A. Bazrafshan, R. L. Bender, A. V. Kellner, H. Ogasawara, R. Ma, F. Schueder, B. G. Petrich, R. Jungmann, R. Li, A.L. Mattheyses, Y. Ke, K. Salaita, Nat. Methods 2020, 17, 1018–1024.

[15] Y. Duan, F. Szlam, Y. Hu, W. Chen, R. Li, Y. Ke, R. Sniecinski, K. Salaita, Nat. Biomed. Eng. 2023, 7, 1404–1418.

[16] M. R. Pawlak, A. T. Smiley, M. P. Ramirez, M. D. Kelly, G. A. Shamsan, S. M. Anderson, B. A. Smeester, D. A. Largaespada, D. J. Odde, W. R. Gordon, Nat. Commun. 2023, 14, 2468.

[17] Y. Zhang, C. Ge, C. Zhu, K. Salaita, Nat. Commun. 2014, 5, 5167.

[18] H. Li, C. Zhang, Y. Hu, P. Liu, F. Sun, W. Chen, X. Zhang, J. Ma, W. Wang, L. Wang, P. Wu, Z. Liu, Nat. Cell Biol. 2021, 23, 642–651.

[19] S. A. Rashid, Y. Dong, H. Ogasawara, M. Vierengel, M. E. Essien, K. Salaita, ACS Appl. Mater. Interfaces 2023, 15, 33362–33372.

[20] Y. Hu, H. Li, C. Zhang, J. Feng, W. Wang, W. Chen, M. Yu, X. Liu, X. Zhang, Z. Liu, Cell 2024, 187, 3445-3459.e15.

[21] R. Merindol, G. Delechiave, L. Heinen, L. H. Catalani, A. Walther, Nat. Commun. 2019, 10, 528.

[22] J. Hahn, S. F. J. Wickham, W. M. Shih, S. D. Perrault, ACS Nano 2014, 8, 8765–8775.

[23] T. Koizumi, Exp. Anim. 1995, 44, 169–171.

[24] T. Koizumi, Exp. Anim. 1995, 44, 181–185.

[25] K. Miyauchi, M. Ogawa, T. Shibata, K. Matsuda, T. Mori, K. Ito, N. Minamiura, T. Yamamoto, Clin. Chim. Acta Int. J. Clin. Chem. 1986, 154, 115–123.

[26] M. Petersen, J. Wengel, Trends Biotechnol. 2003, 21, 74–81.

[27] E. Poppleton, R. Romero, A. Mallya, L. Rovigatti, P. Šulc, Nucleic Acids Res. 2021, 49, W491–W498.

[28] A. Sengar, T. E. Ouldridge, O. Henrich, L. Rovigatti, P. Šulc, Front. Mol. Biosci. 2021, 8, 693710.

[29] M. Pfaff, K. Tangemann, B. Müller, M. Gurrath, G. Müller, H. Kessler, R. Timpl, J. Engel, J. Biol. Chem. 1994, 269, 20233–20238.

[30] B. Ladoux, R.-M. Mège, Nat. Rev. Mol. Cell Biol. 2017, 18, 743–757.

[31] B. Yang, B. Zhou, C. Li, X. Li, Z. Shi, Y. Li, C. Zhu, X. Li, Y. Hua, Y. Pan, J. He, T. Cao, Y. Sun, W. Liu, M. Ge, Y. R. Yang, Y. Dong, D. Liu, Angew. Chem. Int. Ed. 2022, 61, e202202520.

[32] S. Klußmann, A. Nolte, R. Bald, V. A. Erdmann, J. P. Fürste, Nat. Biotechnol. 1996, 14, 1112–1115.

[33] B. E. Young, N. Kundu, J. T. Sczepanski, Chem. Weinh. Bergstr. Ger. 2019, 25, 7981–7990.

[34] W. G. Purschke, D. Eulberg, K. Buchner, S. Vonhoff, S. Klussmann, Proc. Natl. Acad. Sci. 2006, 103, 5173–5178.

[35] S. Helmling, C. Maasch, D. Eulberg, K. Buchner, W. Schröder, C. Lange, S. Vonhoff, B. Wlotzka, M. H. Tschöp, S. Rosewicz, S. Klussmann, Proc. Natl. Acad. Sci. U. S. A. 2004, 101, 13174–13179.

[36] A. Vater, S. Klussmann, Drug Discov. Today 2015, 20, 147–155.

[37] A. M. Kabza, B. E. Young, J. T. Sczepanski, J. Am. Chem. Soc. 2017, 139, 17715–17718.

[38] Z. Wang, W. Xu, L. Liu, T. F. Zhu, Nat. Chem. 2016, 8, 698–704.

[39] A. Pech, J. Achenbach, M. Jahnz, S. Schülzchen, F. Jarosch, F. Bordusa, S. Klussmann, Nucleic Acids Res. 2017, 45, 3997–4005.

[40] Y. Xu, T. F. Zhu, Science 2022, 378, 405–412.

