## Supplementary Information for "Nuclease-Resistant L-DNA Tension Probes Enable Long-Term Force Mapping of Single Cells and Cell Consortia"

[b] Prof. C. Jacob
Department of Biology
University of Mainz
Hanns-Dieter-Hüsch-Weg 15, 55128 Mainz (Germany)

### Experimental Materials and Methods

#### 1. Materials

Cyclo[Arg-Gly-Asp-d-Phe-Lys(PEG-PEG-azide)](RGD-3759-PI) was purchased from biosynth. Neutravidin (31000) (Thermo-Fisher Scientific), Bovine Serum Albumin biotinylated (29130) (Thermo-Fisher Scientific), 25 mm x 75 mm glass coverslips (10812), and sticky slide 8 well (80828) were purchased from ibidi. All oligonucleotides were purchased from Biomers. All buffers were prepared with nuclease free water.

#### 2. General Characterization Methods and Instruments

##### *2.1 TIRF and Brightfield Microscopy*

TIRF and Brightfield Microscopy were performed on a Zeiss Elyra 7 Imaging System equipped with 405nm, 488nm, 561nm, and 642nm excitation lasers using alpha Plan-Apochromat 63x, N.A. 1.46.oil immersion, TIRF objective, pco.edge 4.2 CLHS water-cooled sCMOS cameras.

##### *2.2 DNA Concentrations*

DNA concentrations were determined using a DeNovix-S-06873 (DeNovix OS 0.8.1 v4.1.5) spectrophotometer with a standard value of 33 µg/$OD_{260}$.

##### *2.3 Statistical Analysis*

p values were calculated by performing a t-Test (Two-Sample Assuming Equal Variances) in Microsoft Excel using a built-in data analysis tool pack.

#### 3. Methods

##### *3.1 Surface Preparation*

Surface preparation method was adapted from previously published protocols.[1] Briefly, the glass coverslips (25 x 75 mm) were adhered to the sticky slide 8 well slides. Wells were coated with BSA biotin (100 µg/mL) in nuclease free water overnight at room temperature. Wells were rinsed 3 times with nuclease-free water and incubated with 100 µg/mL of neutravidin for 30 minutes at room temperature. Wells were rinsed once more and incubated at room temperature with 200 µL of 100 nM DNA probes for 1 hour. After washing with nuclease free water, cell culture media was added to the well followed by the addition of cells.

##### *3.2 DNA Hybridization*

DNA oligonucleotides and DNA hairpins were hybridized at 10 µM in 200 µL PCR tubes and subsequently diluted to 100 nM. DNA oligonucleotides were heated to 95 °C and then cooled at a rate of 1.3 °C/min to 25 °C.

##### *3.3 DNA Oligonucleotide Coupling to Cyclic RGD and Purification*

To conjugate cRGDfk to DNA, we used azide/DBCO click chemistry. The DNA was modified with DBCO and the cRGDfk peptide with azide. The reaction was performed in a molar ratio of DNA-DBCO (1mM)/cRGDfk-$N_3$ (3mM) = 1:3 in PBS at 37 °C at 650 rpm for overnight. Excess cRGDfk-$N_3$ was removed using a 3 kDa MWCO spin filter. The product was confirmed with HPLC and MALDI-ToF MS (Supplementary Figure S8).

##### *3.4 Cell Culture*

NIH/3t3 fibroblasts and A10 myoblast cells were cultured according to DMSZ guidelines. Briefly, NIH/3t3 fibroblasts cells were cultured in DMEM supplemented with 10% fetal bovine serum (v/v) and penicillin/streptomycin in an incubator with 5% $CO_2$. A10 cells were cultured in DMEM supplemented with 20% fetal bovine serum (v/v) and penicillin/streptomycin in an incubator with 5% $CO_2$.

#### 4. Supplementary Tables

**Table S1.** DNA sequences for the oligonucleotides with their abbreviations, sequence, and modifications.

| Name | Sequence 5’→ 3’ | Figure | 5’Modification | 3’Modification |
|---|---|---|---|---|
| D-DNA Unzipping Mode | GAG GAG GGC AGC AAA CGG GAA GAG TCT TCC TTT ACG TTT T | Fig. 2c, 4c | ATTO 647N | Biotin |
| D-DNA Ligand Strand | ACG TAA AGG AAG ACT CTT CCC GTT TGC TGC CCT CCT C | Fig. 2c, 4c | DBCO | BHQ 2 |
| L-DNA Unzipping Mode | GAG GAG GGC AGC AAA CGG GAA GAG TCT TCC TTT ACG TTT T | Fig. 2b, 4b | ATTO 647N | Biotin |
| L-DNA Ligand Strand | ACG TAA AGG AAG ACT CTT CCC GTT TGC TGC CCT CCT C | Fig. 2b, 4b | DBCO | BHQ 2 |
| D-DNA Anchor strand with BHQ 2 | CGCATCTGTGCGGTATTTCACTTT | Fig. 3c | BHQ 2 | Biotin |
| D-DNA RGD strand | TTT GCT GGG CTA CGT GGC GCT CTT | Fig. 3c | DBCO | Cy3B |
| D-DNA Hairpin Strand | GTG AAA TAC CGC ACA GAT GCG TTT-GCG CGC GCG CGC TTT TGC GCG CGC GCG C-TTT AAG AGC GCC ACG TAG CCC AGC | Fig. 3c | None | None |
| L-DNA Anchor strand with BHQ 2 | CGCATCTGTGCGGTATTTCACTTT | Fig.3b | BHQ 2 | Biotin |
| L-DNA RGD strand | TTT GCT GGG CTA CGT GGC GCT CTT | Fig. 3b | DBCO | Cy3B |
| L-DNA Hairpin Strand | GTG AAA TAC CGC ACA GAT GCG TTT-GCG CGC GCG CGC TTT TGC GCG CGC GCG C-TTT AAG AGC GCC ACG TAG CCC AGC | Fig.3b | None | None |

#### 5. Supplementary Figures

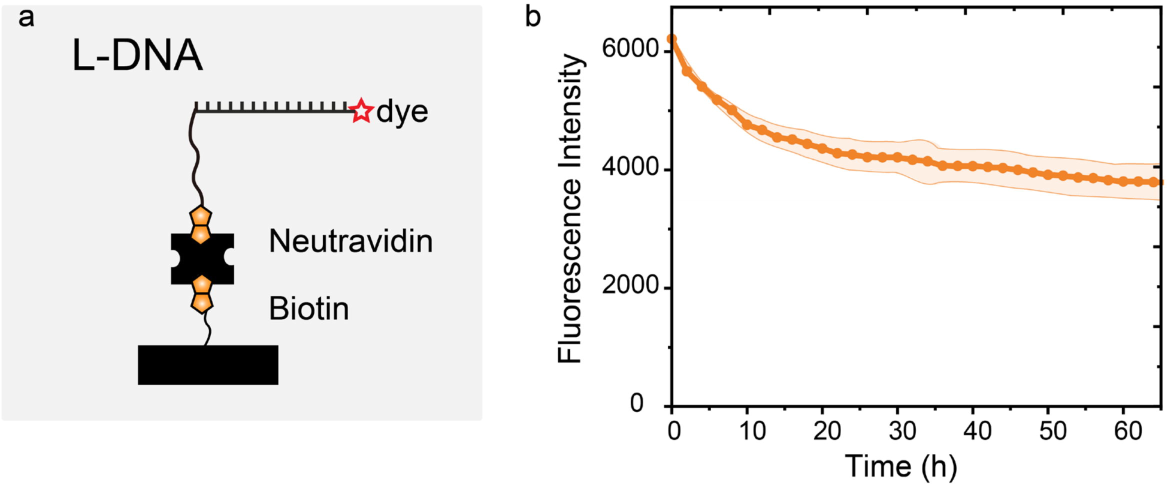

*Figure S1. Assessment of stability of L-DNA immobilized on biotin-neutravidin surfaces in cell culture environment at 37 ˚C. (a) Scheme depicting the immobilization of ss-L-DNA. (b) Mean fluorescence intensity (FI) captured using TIRF illumination over a period of 3 days.*

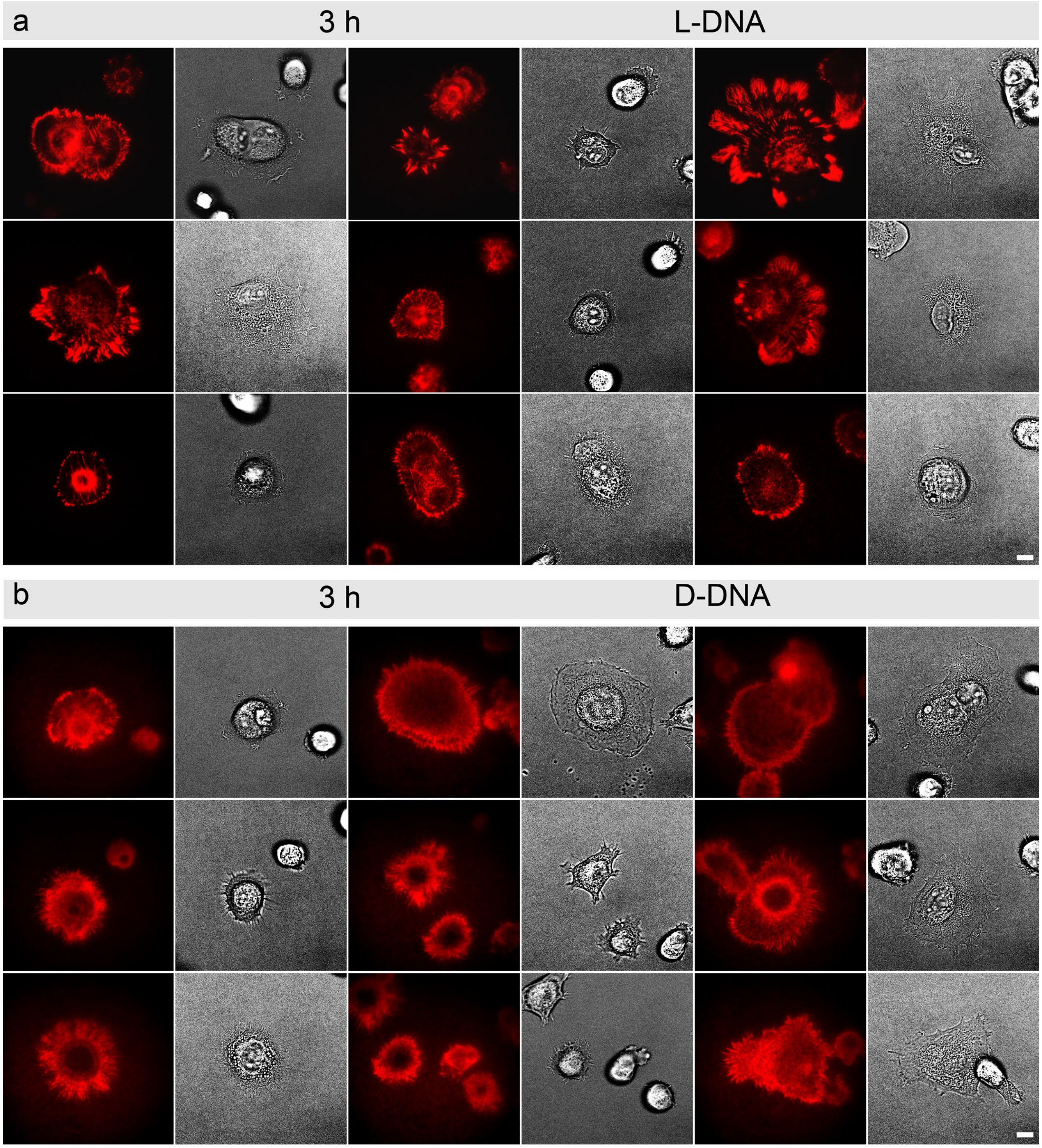

*Figure S2. Representative Images of tension signals made by fibroblasts as they adhere, spread, crawl at 3 h time point. (a-b) Brightfield and fluorescence images on (a) L-DNA and (b) D-DNA tension probe surfaces. Scale bars = 10 μm*

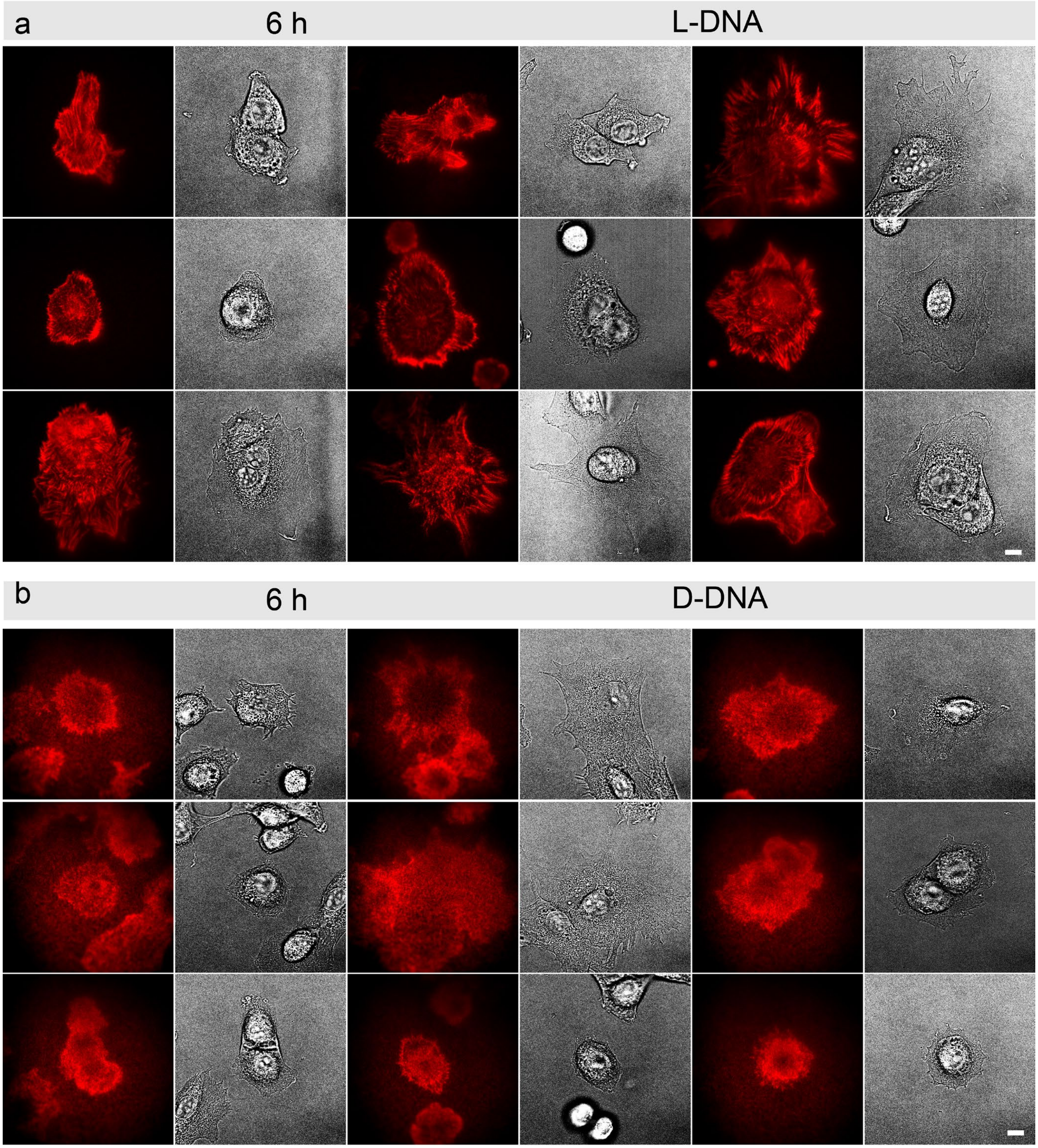

*Figure S3. Representative Images of tension signals made by fibroblasts as they adhere, spread, crawl at 6 h time point. (a-b) Brightfield images and fluorescence signals on (a) L-DNA and (b) D-DNA tension probe surfaces. Scale bars = 10 μm*

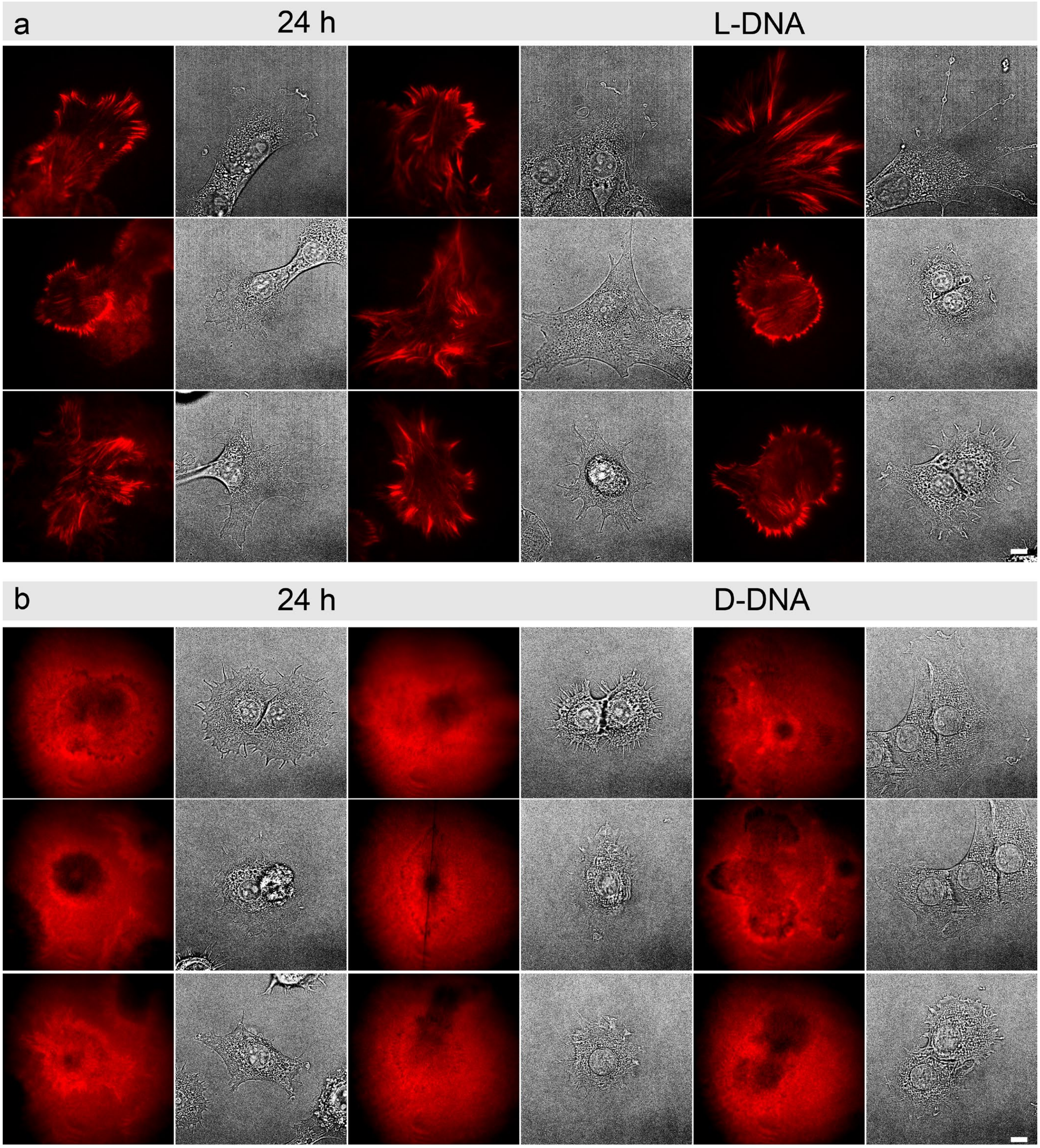

*Figure S4. Representative Images of tension signals made by fibroblasts as they adhere, spread, crawl at 24 h time point (a-b) Brightfield images and fluorescence signals on (a) L-DNA and (b) D-DNA tension probe surfaces. Scale bars = 10 μm*

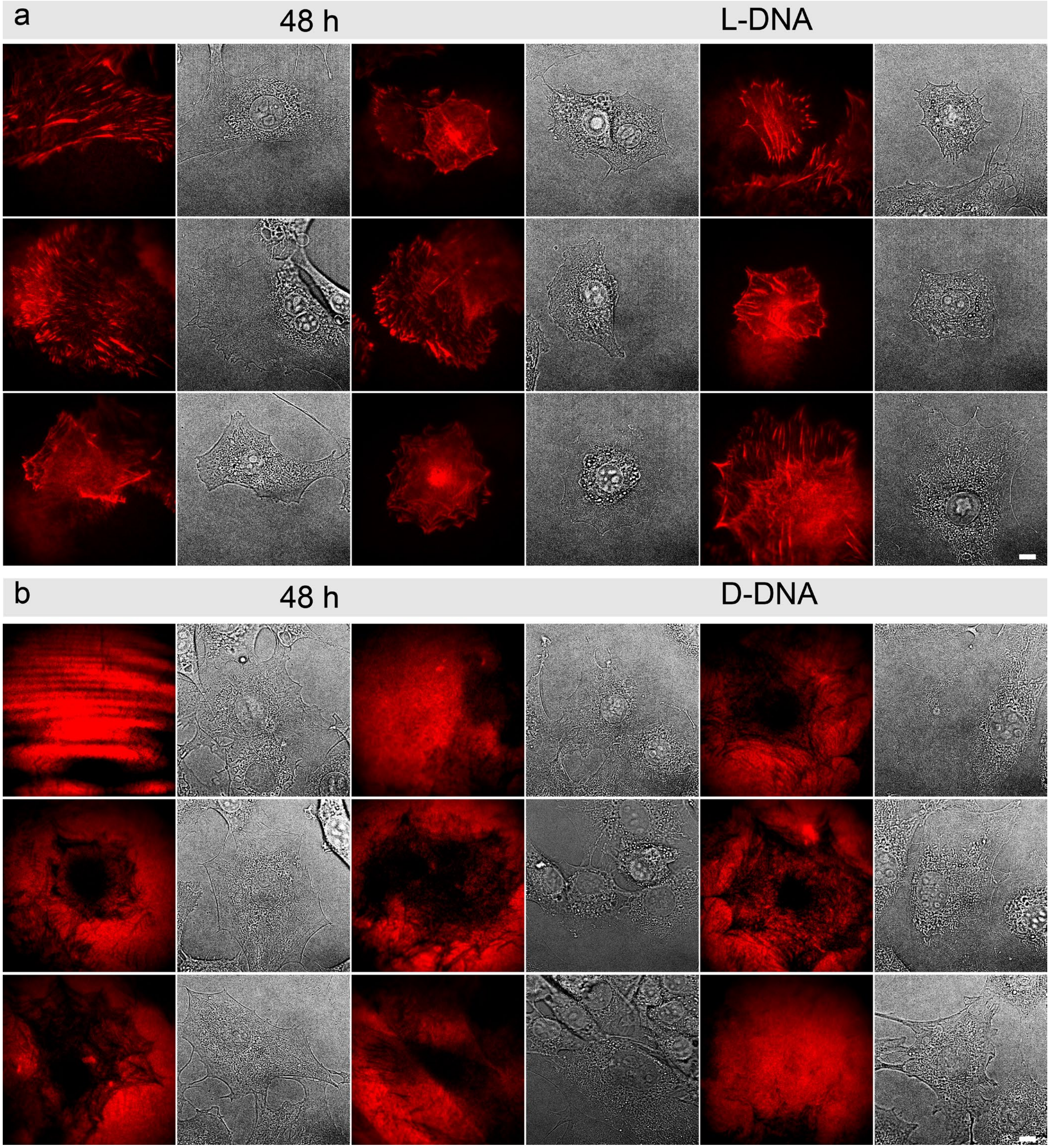

*Figure S5. Representative Images of tension signals made by fibroblasts as they adhere, spread, crawl at 48h time point (a-b) Brightfield images and fluorescence signals on (a) L-DNA and (b) D-DNA tension probe surfaces. Scale bars = 10 μm*

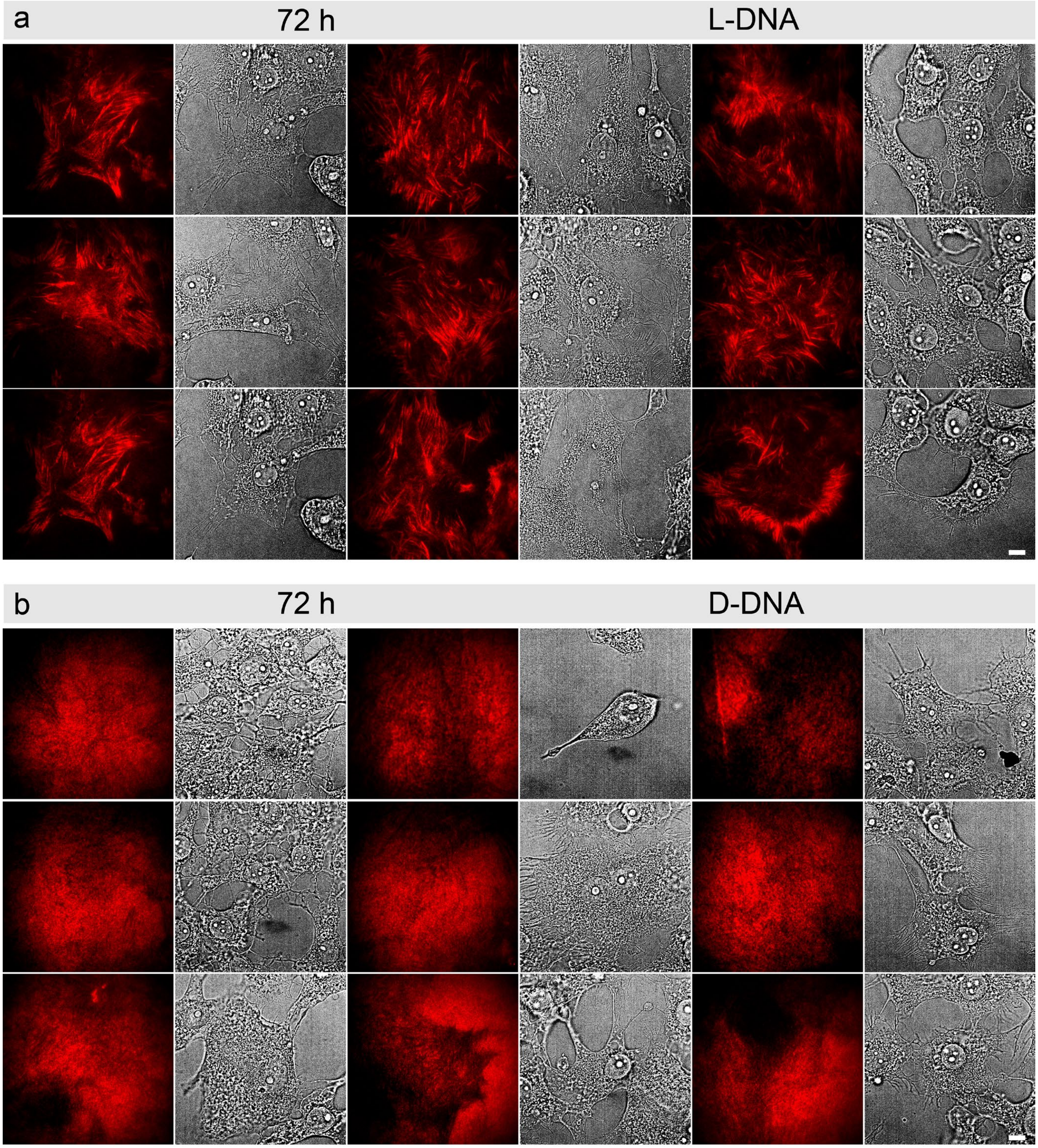

*Figure S6. Representative Images of tension signals made by fibroblasts as they adhere, spread, crawl at 72 h time point (a-b) Brightfield images and fluorescence signals on (a) L-DNA and (b) D-DNA tension probe surfaces. Scale bars = 10 μm*

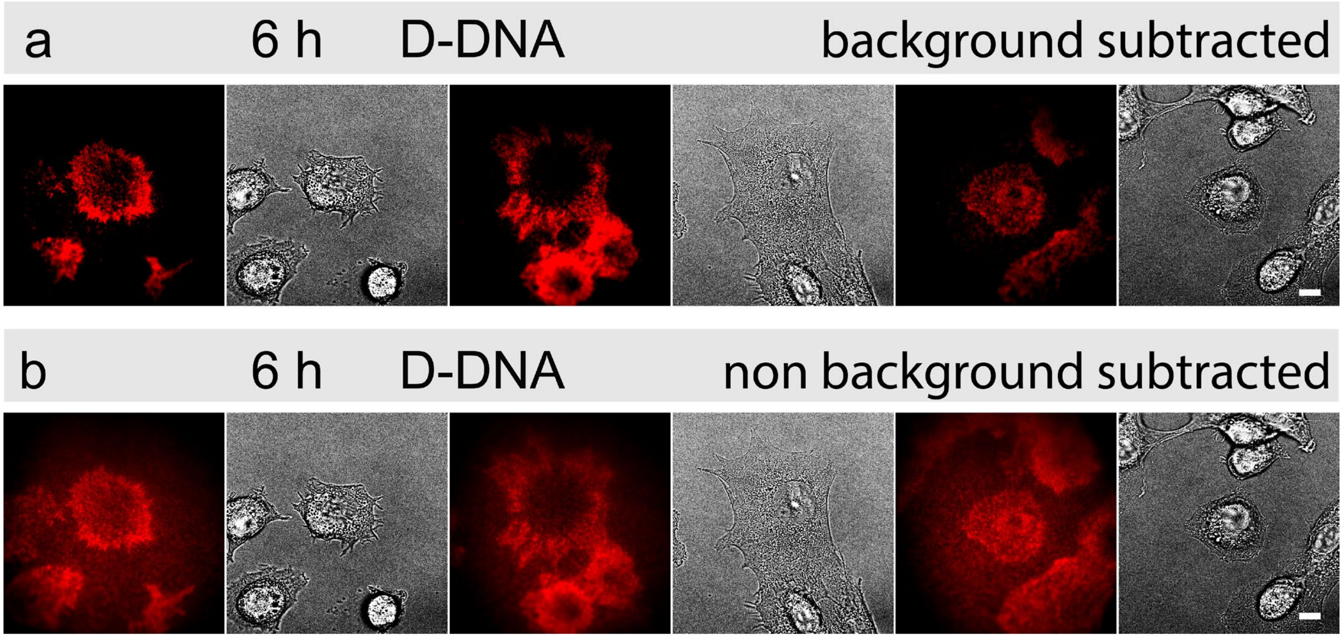

*Figure S7. Representative Images of tension signals at 6 h time point on D-DNA surfaces. (a) Images with routinely used background subtraction (b) Images without any background subtraction. Scale bars = 10 µm*

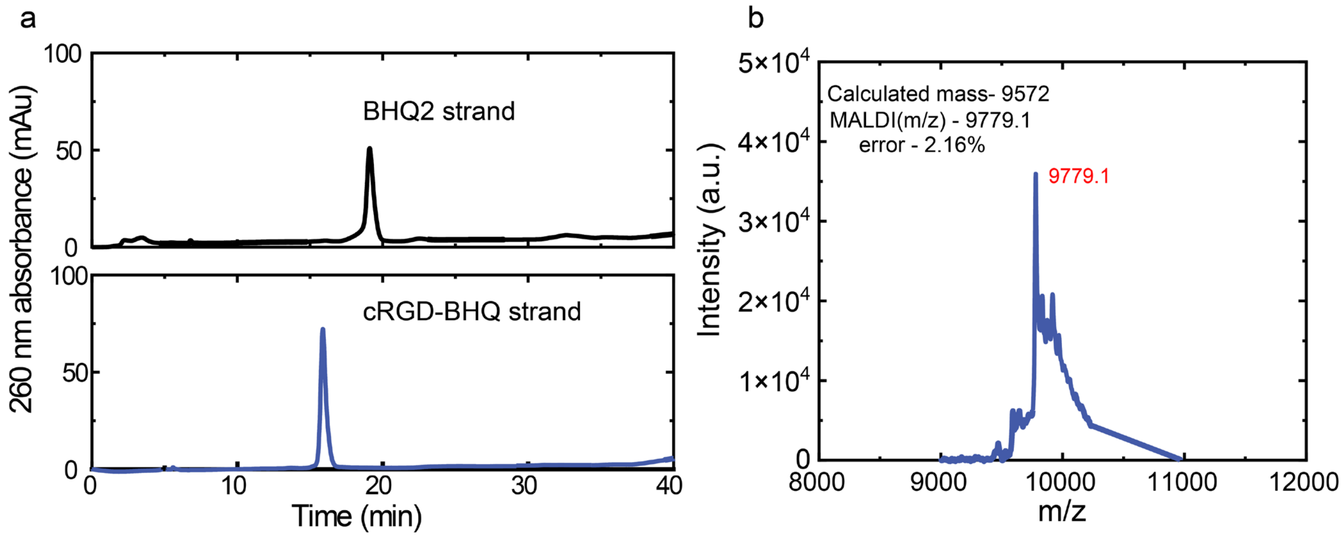

*Figure S8. Characterization of DNA conjugated with cRGD. (a) HPLC graphs depicting different elution times of DNA-BHQ ssDNA and cRGD-DNA-BHQ ssDNA. (b) MALDI-ToF MS depicting the observed molecular weight of the cRGD-DNA-BHQ ssDNA product. Calculated for [M+9Na] + = 9779, Found [M+9Na] + = 9779.1.*

#### 6. Supplementary References

[1] M. R. Pawlak, A. T. Smiley, M. P. Ramirez, M. D. Kelly, G. A. Shamsan, S. M. Anderson, B. A. Smeester, D. A. Largaespada, D. J. Odde, W. R. Gordon, *Nat. Commun.* **2023**, *14*, 2468.
